# The anaerobic digestion microbiome: a collection of 1600 metagenome-assembled genomes shows high species diversity related to methane production

**DOI:** 10.1101/680553

**Authors:** Stefano Campanaro, Laura Treu, Luis M Rodriguez-R, Adam Kovalovszki, Ryan M Ziels, Irena Maus, Xinyu Zhu, Panagiotis G. Kougias, Arianna Basile, Gang Luo, Andreas Schlüter, Konstantinos T. Konstantinidis, Irini Angelidaki

## Abstract

**Background:** Microorganisms in biogas reactors are essential for degradation of organic matter and methane production through anaerobic digestion process. However, a comprehensive genome-centric comparison, including relevant metadata for each sample, is still needed to identify the globally distributed biogas community members and serve as a reliable repository.

**Results:** Here, 134 publicly available datasets derived from different biogas reactors were used to recover 1,635 metagenome-assembled genomes (MAGs) representing different bacterial and archaeal species. All genomes were estimated to be >50% complete and nearly half were ≥90% complete with ≤5% contamination. In most samples, specialized microbial communities were established, while only a few taxa were widespread among the different reactor systems. Metabolic reconstruction of the MAGs enabled the prediction of functional traits related to biomass degradation and methane production from waste biomass. An extensive evaluation of the replication index provided an estimation of the growth rate for microbes involved in different steps of the food chain. The recovery of many MAGs belonging to Candidate Phyla Radiation and other underexplored taxa suggests their specific involvement in the anaerobic degradation of organic matter.

**Conclusions:** The outcome of this study highlights a high flexibility of the biogas microbiome. The dynamic composition and adaptability to the environmental conditions, including temperatures and a wide range of substrates, were demonstrated. Our findings enhance the mechanistic understanding of anaerobic digestion microbiome and substantially extend the existing repository of genomes. The established database represents a relevant resource for future studies related to this engineered ecosystem.

## Background

Anaerobic environments are ubiquitous in the biosphere, some examples are the digestive tract of animals, paddy fields and aquatic sediments. These environments play crucial roles in the degradation of organic matter and in the global carbon cycle [1,2]. The anaerobic digestion (AD) process is also an example of such an environment, having great societal importance since it reduces our dependence on fossil fuels via its ability to generate methane within engineered bioreactors [3]. For these reasons, the AD process has been widely acknowledged as an efficient biochemical route allowing the conversion of organic wastes into energy and other valuable products, and has been posited as a sustainable solution for resource recovery and renewable energy production underpinning the circular economy concept [4]. Apart from biowaste valorization, this process is also of great importance for nutrient recycling, since it stabilizes the nitrogen compounds making them more easily assimilable by plants, reducing nitrogen leakage to soil and groundwater.

Methane is one of the most relevant end-products generated during the methanogenesis step of AD process, and is produced by methanogenic *Archaea* [5]. While in the rumen methanogenesis results in loss of energy and carbon to the atmosphere [6], in anaerobic digesters methane production and purity is maximized for exploiting the economic value of the process. Methane production has been directly linked to the composition of the AD microbiome [7–9], and it is also under the control of microbial metabolism, which is in turn thermodynamically dependent on environmental parameters of the reactor [10,11]. The intimate connection between these parameters offers unique opportunities to improve process efficiency, which can be obtained through microbial selection or manipulation [12,13].

In the past four years, hundreds of genomes belonging to the microbes involved in biogas production process have been sequenced. To improve the understanding of the highly diverse and interconnected network of the AD microbiome, several studies focused on the characterization of microbial communities from the full-scale biogas plants [14–17] trying to connect the microbial composition to differences in process parameters [7,18]. Other studies were focused on the identification of the functional roles of some microbial species [19–22]. More frequently, laboratory-scale reactors were studied, since they offer the unique opportunity to implement different configurations, modulate individual process parameters, or explore different feedstock compositions and inocula [23–30]. Additionally, the use of stable isotope labelling experiments on samples collected from lab-scale reactors allowed for the identification of specific substrate utilization abilities by different microbes [31,32]. Despite the great value of these previous studies, the entire datasets of microbial genomes identified have not been previously compared or combined, limiting our global perspective of the AD microbiome.

Cultivation-based approaches to isolate microorganisms from the AD environment have yielded hundreds of novel species and strains (see for example Maus *et al*., 2017). However, this approach is limited to the cultivable fraction of the community, which was estimated between 20% and 95% for deeply studied anaerobic environments such as human gut [33]. To get insights into the genetic repertoire of non-cultivable biogas community members, metagenome sequencing, including assembly and binning strategies, has been highly valuable. Corresponding research has led to the compilation of MAGs frequently representing novel taxa whose lifestyle could be hypothesized based on nucleotide sequence information, thus, making this approach popular recently. Genome-centric metagenomic approaches have been leveraged to obtain large numbers of MAGs across many different environments, including -for example-913 MAGs collected from the cow rumen [34], 957 from the TARA Oceans metagenomes [35], 2,540 from sediment-hosted aquifers [36], and almost 8,000 from a range of environments covered by public sequence data [37]. However a similar approach has not yet been applied to AD systems alone, and therefore, there is a need for a study which would disclose the “AD black box” and assign roles to the species involved in the food chain of the organic matter degradation process.

To fulfill the need of a large scale AD microbial genome database, we present a comprehensive metagenome-centric analysis performed by incorporating nearly 0.9 Tbp of sequence data currently available in the public databases, representing a wide range of different biogas reactor systems. The use of a homogeneous assembly and binning workflow, associated with a de-replication strategy, identified the genomes of nearly 1,600 distinct bacterial and archaeal species. The resource presented in this study provides an important reference dataset for understanding the relative composition and function of the AD microbiome associated with different reactors systems and process parameters. Phylogenetic, biochemical and bioinformatics performance results were investigated in order to reveal a microbiota-functionality nexus that can assist in the development of a more efficient management strategy for the AD system.

## Results and discussion

### Public metagenomes selection and data processing

To get an overview of the AD microbiome, 18 experiments published within 2018 were selected. These include 134 samples, some of them representing biological replicates (Fig. 1). Among these datasets, both laboratory-scale- and full-scale biogas plants fed with a range of different substrates were considered. Details regarding the specific reactor conditions, feedstock and biochemical parameters are reported in Additional File 1 along with other metadata. Most samples were collected from laboratory-scale biogas reactors and batch tests, while other samples were obtained from 23 full-scale biogas plants located in Europe (marked with “*” in figure 1; two of them were sampled at different time points).

**Fig. 1.**
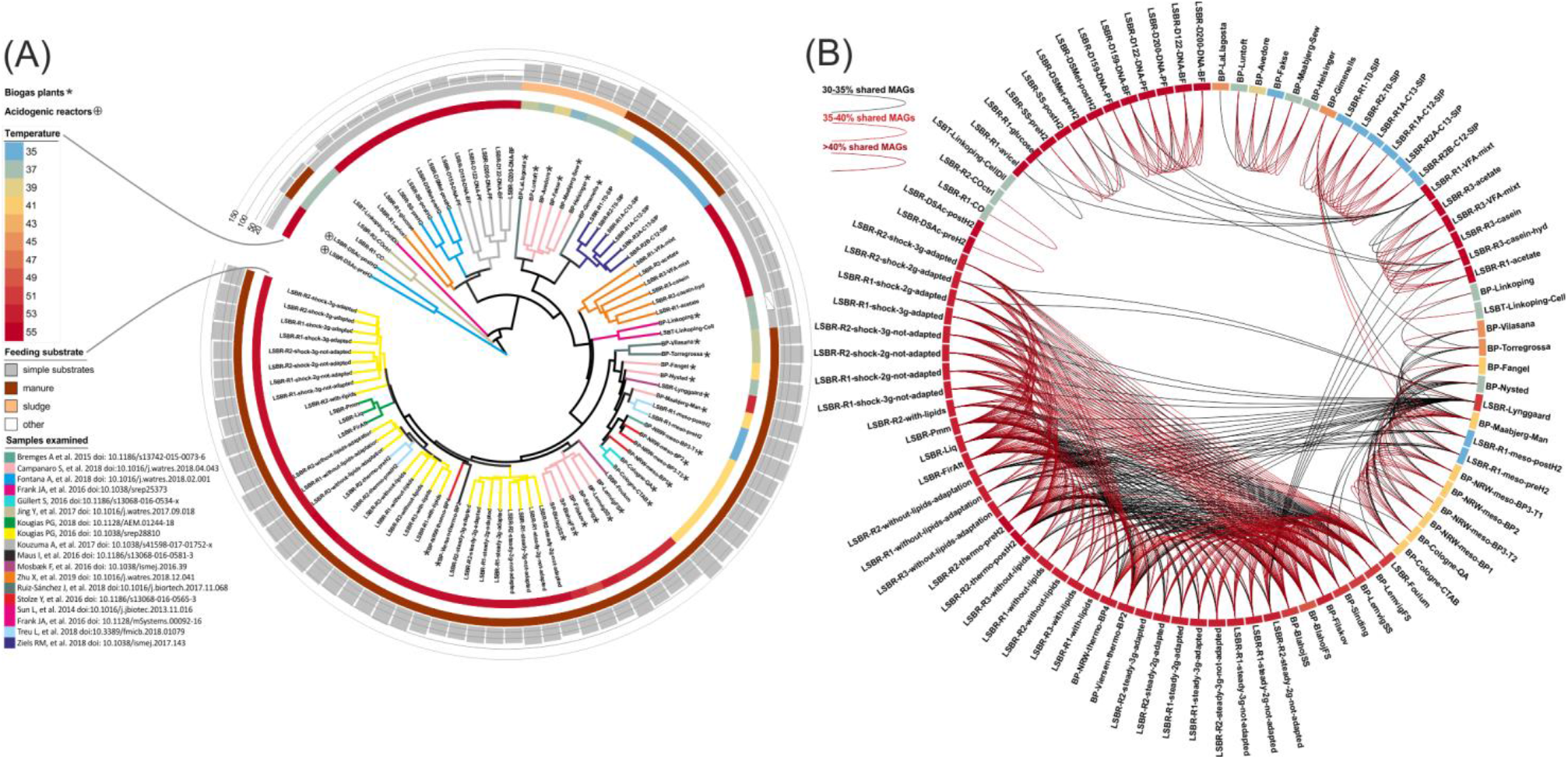
(A) The tree is a representation of the β-diversity values determined comparing the samples. Colors in the tree refer to different experiments reported in the inset (down-left). Reactors temperature and feeding substrates are reported in the external circles. Histogram graph in the external ring represents Fisher alpha diversity values. Samples marked with asterisks were collected from full-scale biogas plants. Details regarding the characteristics of samples are reported in Additional file 1. (B) Representation of the fraction of MAGs “shared” among samples. Arcs colored from black to dark red connect samples having increasing fractions of shared MAGs. Samples in the external circle are colored according to the temperature of the reactor and in (A) and (B) are reported in the same order.

Microbial composition was initially determined considering unassembled reads, and this highlighted marked differences between samples, which were classified into 35 groups. This microbial diversity is also clearly evident in figure1 B, where different samples are connected with arcs having different colors depending on the fraction of common species. The high diversity between groups indicated the need for a separate assembly of each group in order to minimize computational requests, as well as to avoid co-assembly of different strains belonging to the same species, a process resulting in lower quality of the assembled MAGs [38]. A subsequent binning approach was independently performed on each assembly of the 35 groups, leading to a total of 5,194 MAGs (Table 1). Data regarding metagenomic assemblies and number of MAGs collected from the binning process are reported in detail in Additional File 2. Those MAGs featuring completenesses (Cp) lower than 50% and/or contamination rates (Ct) higher than 10% were discarded from the dataset. The remaining MAGs were de-replicated by means of the genome-aggregate Average Nucleotide Identity (ANI) value, in this case standing for species recognition (Additional File 3) [39]. The de-replication step reduced the number of MAGs down to 1,635 unique ones, which represent the collection of bacterial and archaeal genomes established in this study (Table 1). According to the metrics recently proposed for evaluation of Cp and Ct of MAG [40], 796 (~49%) were of high quality (HQ), the remaining were defined as medium-high quality (MHQ) and medium quality (MQ) (Table 1; Fig. 2). By considering all 134 samples, on average 89% of the reads were consistently aligned on the 1,635 MAGs, suggesting that the obtained dataset captured much of the available sequencing information. Results obtained were quite similar when only the HQ MAGs were selected. The degree of novelty of our study was determined performing a comparison with MAGs previously recovered from the AD environment [23,24,31] (https://biogasmicrobiome.com/). Our study showed an improvement in the quality (increased Cp and/or reduced Ct) of 75% of the MAGs already present in public repositories, and added 1228 “new species”, consistently improving the entire biogas microbiome (Additional File 4).

**Table 1.** Number of MAGs assigned to different categories according to their quality. MAGs belonging to one cluster generated during ANI calculation are indicated as “Selected from clusters*”, while MAGs not clustered at more than 95% ANI are indicated as “identified once**”. (ANI=genome-wide Average Nucleotide Identity).

|  | MAGs number<br>before clustering (ANI) |  | MAGs number<br>after clustering (ANI) |
| --- | --- | --- | --- |
| HQ MAGs<br>Cp>90%, Ct<5% | 1628 | HQ MAGs selected<br>from clusters* | 441 |
|  |  | HQ MAGs identified<br>once | 355 |
| MHQ MAGs<br>90%>Cp>=70%,<br>10%>Ct>5% | 1316 | MHQ MAGs selected<br>from clusters | 170 |
|  |  | MHQ MAGs identified<br>once | 435 |
| MQ MAGs<br>50%>=Cp>70%,<br>10%>Ct>5% | 526 | MQ MAGs selected<br>from clusters | 15 |
|  |  | MQ MAGs identified<br>once | 219 |
| LQ MAGs<br>Cp<50% | 1436 |  |  |
| Contaminated<br>MAGs<br>Ct>10% | 288 |  |  |
| Total | 5194 | Total | 1635 |

**Fig. 2.**
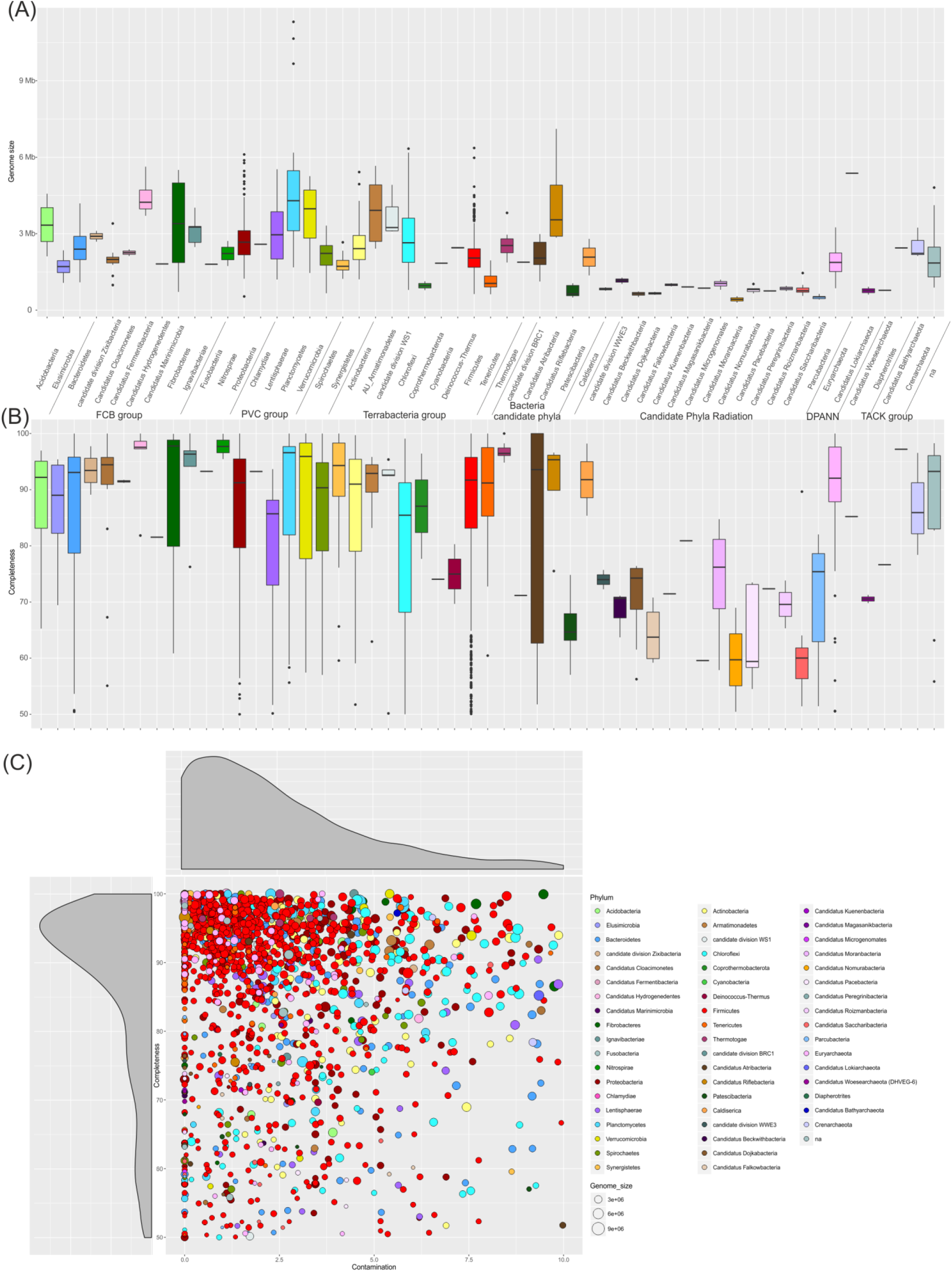
Box plots reporting (A) genome size and (B) completeness of the 1,635 selected MAGs. (C) Scatter plot reporting the completeness and contamination levels for each MAG. Colors refer to the taxonomic assignment at phylum level and in (C) circle size is proportional to the genome length.

### Structure of the microbial community

Analyses performed using the Microbial Genomes Atlas (MiGA) Online [41] estimated that a relevant fraction of the genomes belong to taxonomic groups for which genomes of type material are not present in the NCBI genome database. More specifically, 0.2% of MAGs cannot be assigned to known phyla, 11.6% to known classes, 69.7% to orders, 71.3% to families, 92.1% to genera and 95.2% to species. This evidenced that the present genome-centric investigation allowed to fill-in a notable gap in the knowledge of the AD microbial community. A dedicated project was established to allow the recovery of both genome sequences of MAGs and their taxonomic assignment “http://microbial-genomes.org/projects/biogasmicrobiome”. Using the AAI approach as implemented in MiGA Online [41], it is also possible to perform queries with genomes of interest using the MAGs identified in the present study as a reference.

In addition to the analysis performed with MiGA, to determine the taxonomic position of the MAGs, a procedure based on four different evidences was used. (1) 1,233 MAGs were taxonomically assigned using selected marker genes and (2) an additional 212 MAG were characterized based on results obtained from 16S rRNA gene sequences. (3) The taxonomy of the 121 remaining MAGs (mainly belonging to candidate taxa) has been refined by manual inspection of their placement into a phylogenetic tree as previously described [7]. (4) Only 69 out of 1,635 MAGs were assigned to known species based on ANI comparison performed considering the genomes deposited in NCBI (https://www.ncbi.nlm.nih.gov/genome/microbes/) (Additional File 3). Furthermore, the vast majority of MAGs (1,574) obtained were assigned to the domain *Bacteria* and only 61 to *Archaea*, and distributed over 55 different phyla as reported in figure 3. However, according to a previous study [42], the vast majority of species were classified as belonging to the phylum *Firmicutes* (790 MAGs), followed by *Proteobacteria* (137 MAGs) and *Bacteroidetes* (126 MAGs). The *Euryarchaeota* (53 MAGs) was the largest archaeal phylum in this study. Numerous species associated to candidate phyla were also identified by our study and were tentatively assigned to *Cloacimonetes* (15 MAGs), *Atribacteria* (8 MAGs), *Saccharibacteria* (8 MAGs) and *Dojkabacteria* (8 MAGs). As evidenced in figure 2 and figure 3, there are marked differences in genome size among MAGs associated to different phyla. For example, *Planctomycetes* and *Chloroflexi* have on average larger genomes (p<0.001), while *Tenericutes* (p<0.001) and CPR taxa, such as Candidatus *Dojkabacteria* (p=0.022), have smaller genomes. Taxonomic assignment was compared with that obtained from MiGA [41] and results obtained were in good agreement; the fraction of MAGs consistently assigned to already existing taxa varied from 68% (family) to 88% (genus) depending on the taxonomic level.

**Fig. 3.**
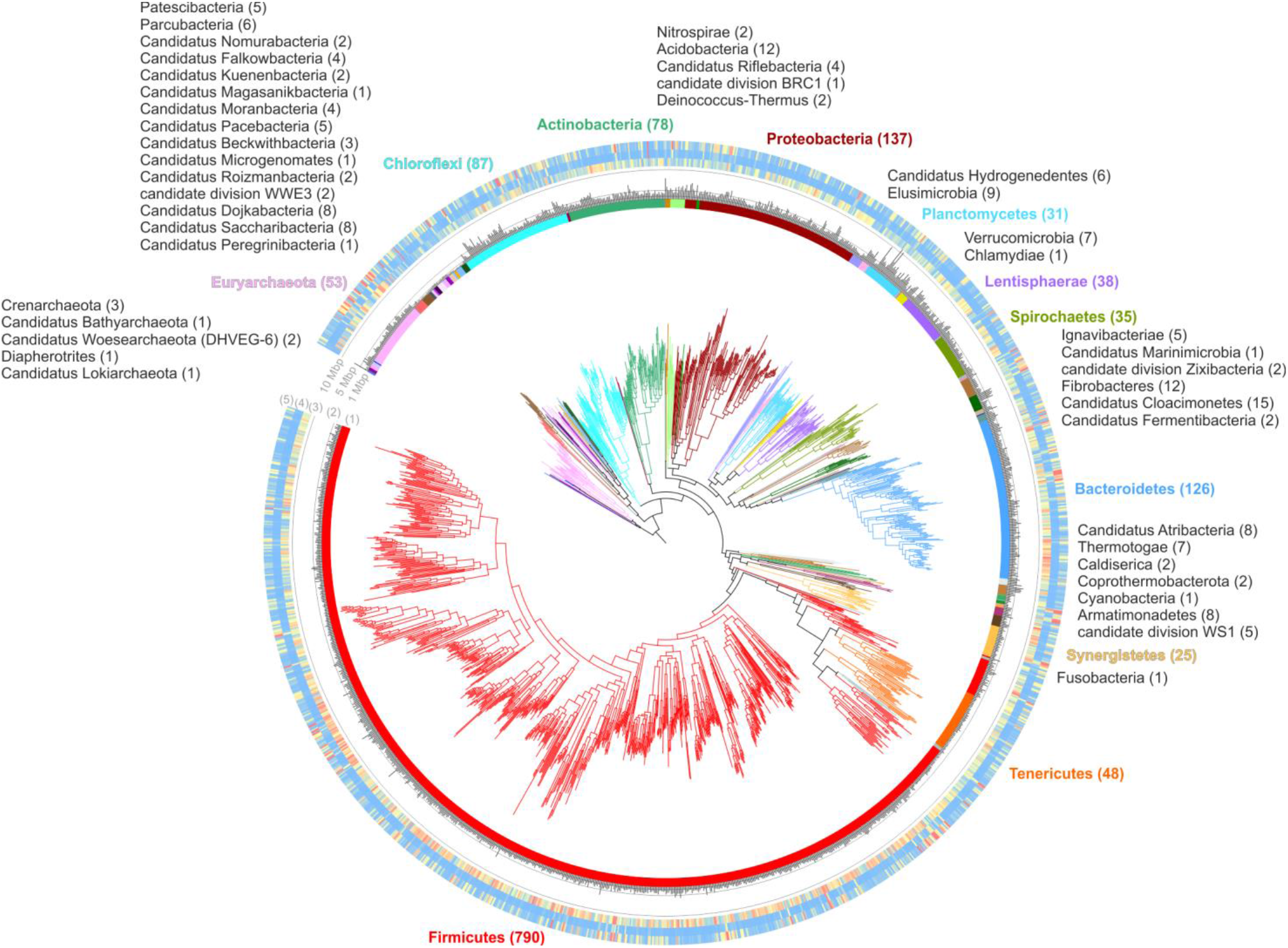
MAGs taxonomic assignment across 54 phyla. The maximum likelihood tree was inferred from the concatenation of 400 taxonomic informative proteins and spans a de-replicated set of 61 archaeal and 1,574 bacterial MAGs. External circles represent, respectively: (1) taxonomic assignment at phylum level, (2) genome size (represented as a bar plot), (3) heatmap representing the number of experiments where each MAG showed the abundance higher than 0.001% (from blue 0% to red 10%), (4) average abundance (from blue 0% to red 10%) and (5) maximum abundance determined among the entire set of experiments (from blue 0% to red 10%).

The reactor process conditions influence the relative abundance of taxa, determining dramatic changes. For instance, the “*Bacteria/Archaea*” ratio, which has a median value of ~14, was highly variable. Beside the acidogenic reactors, where the methanogenic process was undetectable (i.e. “LSBR-DSAc-preH2” and “LSBR-DSAc-postH2”), it was concluded that in 7.7% of all samples archaeal abundance was lower than 1% and consequently “*Bacteria/Archaea*” ratio exceeded 100. However, *Archaea* were predominant in several reactors analyzed in this study and in 3% of all samples, their abundance exceeded that of *Bacteria*, with a ratio of ~0.5 in a biofilm sample collected from a reactor fed with acetate (“LSBR-D200-DNA-BF”). Despite the fact that some of the microbiome derived from sub-fractions of the samples (e.g. stable isotope labelling) or from biofilms, it is interesting to note that in reactors fed only with “methanogenic substrates”, *Archaea* was the dominant group of the entire microbiome. Considering only biogas plants, the “*Bacteria/Archaea*” ratio is kept within a more narrow range, but still it is very flexible (from 470 in Nysted to 3.4 in Vilasana) (Fig. 4). The bacterial phylum *Firmicutes*, which is the most abundant taxon within the biogas microbiome, also varied between 1.3% and 99.9% of the microbial community (Additional Fig. S1 and Additional File 5). In almost 40% of all samples analyzed, *Firmicutes* was not the dominant taxon, but *Bacteroidetes, Coprothermobacter, Actinobacteria, Thermotogae, Chloroflexi* and *Euryarchaeota* become prevalent reaching up to 85% relative abundance within the microbiome [42]. Interestingly, in reactors where none of the previously mentioned taxa were dominant, microbial species belonging to candidate phyla reached high relative abundances, as was the case for *Candidatus* Cloacimonetes (15.7%), *Ca*. Fermentibacteria (16.4%), *Ca*. Roizmanbacteria (19%) and *Ca*. Saccharibacteria (16.4%) (Additional File 5). High abundance of *Ca*. Fermentibacteria in full-scale mesophilic anaerobic digesters at Danish wastewater treatment facilities was previously reported [43]. These findings evidence that the composition of the AD microbiome has a remarkable flexibility. Additionally, the high relative abundance of yet-uncultivated taxa suggests that they may play an important role in the microbial community, and probably warrant further attention.

**Fig. 4.**
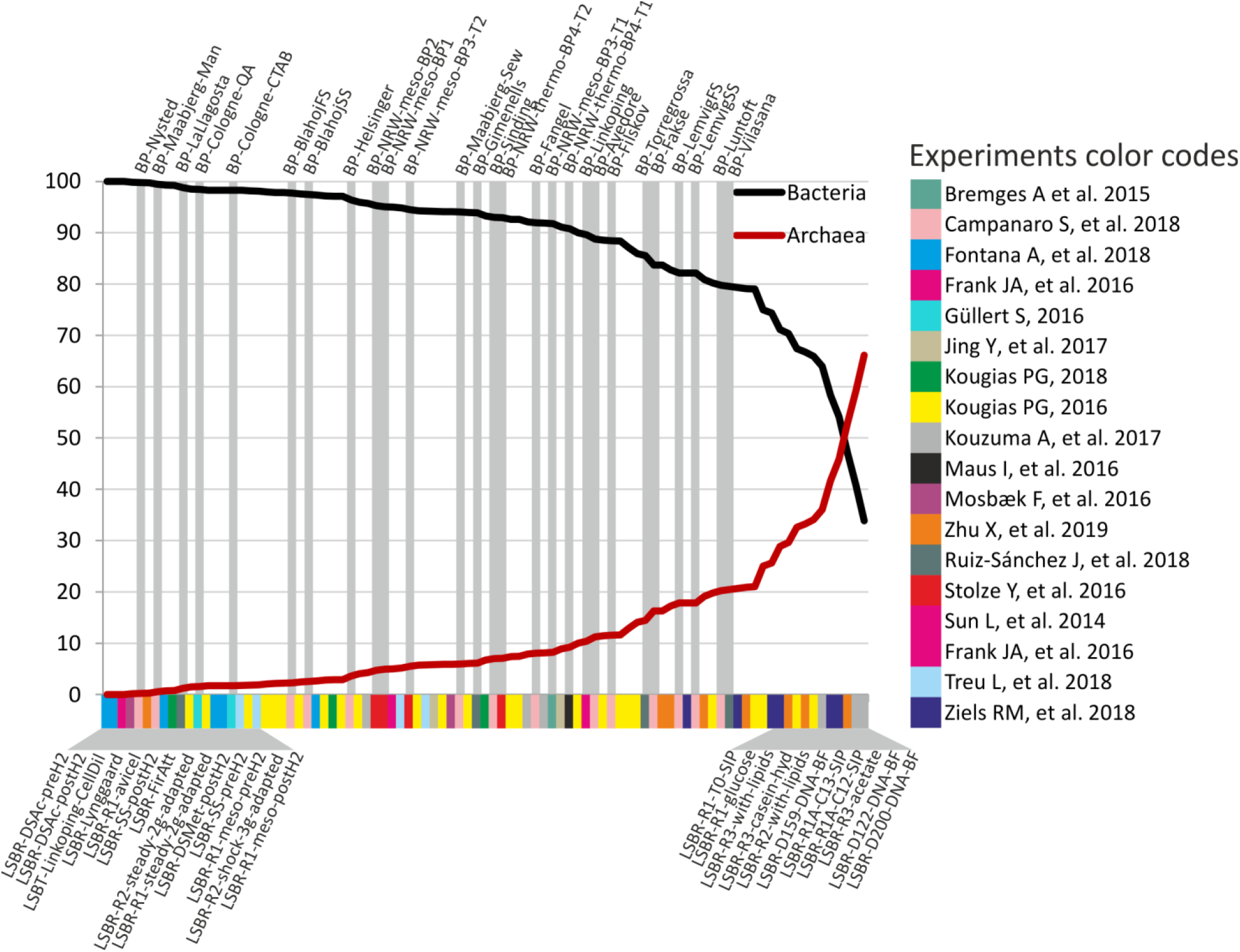
Coverage values of all the MAGs calculated in each sample were used to estimate relative abundance of *Archaea* and *Bacteria*. Gray stripes in figure highlight samples collected from biogas plants, while at the bottom part of the figure color codes were used to distinguish the 18 “experiments” investigated (see inset in the right part of the figure).

We calculated the variation in abundance for each MAG across the AD samples, along with their taxonomic assignment. Additionally, the number of MAGs in each sample was estimated considering as “present” those with abundances higher than 0.001%. This analysis revealed that the microbial community composition was highly variable depending on the AD sample, as clearly reported in figures 1 A and B and Additional Fig. S2. The number of detectable species in the microbiome ranged between 79 (Fisher alpha diversity 4.4) and 1213 (Fisher alpha diversity 133.8) (Additional File 6). This wide range was due to the high variability of the environmental parameters and feedstock compositions. According to previous findings [8,28], thermophilic reactors have a lower number of species than mesophilic (p-value<0.001). Among the thermophilic reactors in this study, those characterized by a very high number of species were fed with manure or a mixture of manure and agricultural feedstocks, while those having few species were fed with simplified substrates such as cheese whey, acetate or glucose (p-value<0.001). This suggests that the AD process can be supported by less than 100 species when the feedstock is mainly consisting of one single compound. On the contrary, degradation of complex substrates (such as sewage sludge or manure) requires the cooperation of a large cohort of microbes including more than 1,000 species. Analysis of the MAGs shared among different samples (Fig. 1 B) revealed that thermophilic reactors tend to share more species than mesophilic systems, which could be due to the selective pressure imposed by the high growth temperature.

Cluster analysis was performed both at individual MAG abundance level and at sample level (rows and columns in Additional Fig. S2) in order to verify MAGs and samples having similar abundance profiles, respectively. This allowed the assignment of MAGs to two main groups: “G1” includes mostly *Chloroflexi* and *Bacteroidetes*, while “G2” includes mostly *Firmicutes*. Sample clustering revealed three main groups, “C1” including reactors fed with sewage sludge, “C2” those fed with “simplified substrates” and “C3” fed with manure only. A similar classification is shown in figure 1, indicating that the feeding substrate is one of the main driving forces of the AD microbiome diversification. A similar classification is shown in figure 1, indicating that the feding substrate is one of the main driving forces of the AD microbiome diversification. Furthermore, the Principal Coordinates Analysis (PCoA) performed considering the microbiome composition originating from different AD environments revealed no clear separation of samples within the plot representing thermophilic and mesophilic reactors (Additional Fig. S3A). This is in contrast to previous findings [5,7] showing mostly specialized microbial communities depending on the temperature regime. We assume that, for most of the species considered in this study, rather the substrate used plays a major role in determining their abundance than the temperature (Additional Fig. S3 B-C). We are aware of the fact that some MAGs did not pass the selection determined by assembly and subsequent binning processes, and this might introduce a shift into the PCoA. However, the MAGs collectively captured on average 89%, and sometimes close to 100%, of the total reads of the metagenomic datasets and thus, the majority of the sampled microbial communities. The collection of MAGs considered in this study does not completely disclose the “AD black box”. However, we are getting closer to the assignment of a substantial number of new species to their role in the food chain of the organic matter degradation process.

Only few MAGs were detected in multiple samples, and this was due to the high heterogeneity of the AD microbiome (Additional Fig. S2; Additional File 7). By considering the highly abundant MAGs (more than 1% relative abundance), only 25 were present in more than 10% of the samples, while 1246 were considered as low abundant (lower than 1%) (Additional Fig. S4). Among the 25 abundant MAGs, four methanogenic *Archaea* were identified, namely the *Candidatus* Methanoculleus thermohydrogenotrophicum AS20ysBPTH_159, *Methanosarcina thermophila* AS02xzSISU_89, *Methanothrix soehngenii* AS27yjCOA_157 and *Methanoculleus thermophilus* AS20ysBPTH_14. The remaining 21 MAGs were assigned to the phyla *Firmicutes* (14 MAGs), *Bacteroidetes* (2 MAGs), *Synergistetes* (2 MAGs), *Thermotogae* (1 MAG) and *Coprothermobacterota* (1 MAG). In general, *Euryarchaeota* were statistically over-represented among the highly abundant MAGs (p<0.05) and this result was clearly due to the origin of the examined samples as they were obtained from AD systems producing methane. Interestingly, *Defluviitoga tunisiensis* AS05jafATM_34, one out of seven MAGs of the phylum *Thermotogae* identified in this study, was present at high abundance (average 2.1%; maximum 58.9%). Widespread identification of this species in reactors suggests its central role in thermophilic AD system. Specific metabolic potential related to saccharide, polyol, lipid transport systems (Additional File 8) and hydrogen production characterize the *Thermotogae* species and possibly explain their importance in AD (Maus et al., 2016a). Analysis of the low abundant MAGs (threshold 0.001%), revealed that 94% of these taxa were present in more than 10% of the samples, and the phyla statistically over-represented in this group were *Chloroflexi, Elusimicrobia, Firmicutes* and *Plantomycetes* (p<0.01). This finding indicates that many MAGs are widespread in the global AD microbiome, but they are present at very low relative abundances. We assume that, under certain favorable conditions, their abundance may increase, and they can become dominant.

### Functional analysis based on KEGG annotation

Metabolic pathway reconstruction and biological role interpretation of 1401 HQ and MHQ MAGs were performed applying a collection of functional units, called KEGG modules. Analysis was performed on 610 modules (present in prokaryotes/KEGG DB), and identified that 76.2% of them are “complete” in at least one MAG, 10.1% have at best one block missing (1 bm) and 2.5% have at best two blocks missing (2 bm). In the following, only complete and “1 bm” modules will be considered. Modules distribution and completeness indicated that a very low number of them is widespread in MAGs, while the majority have a scattered distribution in terms of presence/absence (Fig. 5). Additionally, the association of many modules with some specific taxa is remarkable; in fact, a strong correlation between the clustering based on modules presence/absence and MAGs taxonomic assignment was found (Fig. 5; Additional File 9). In particular, only 15 modules are present in more than 90% of the phyla (widespread), 108 are present in 50-90% (frequent) and 434 in less than 50% (infrequent).

**Fig. 5.**
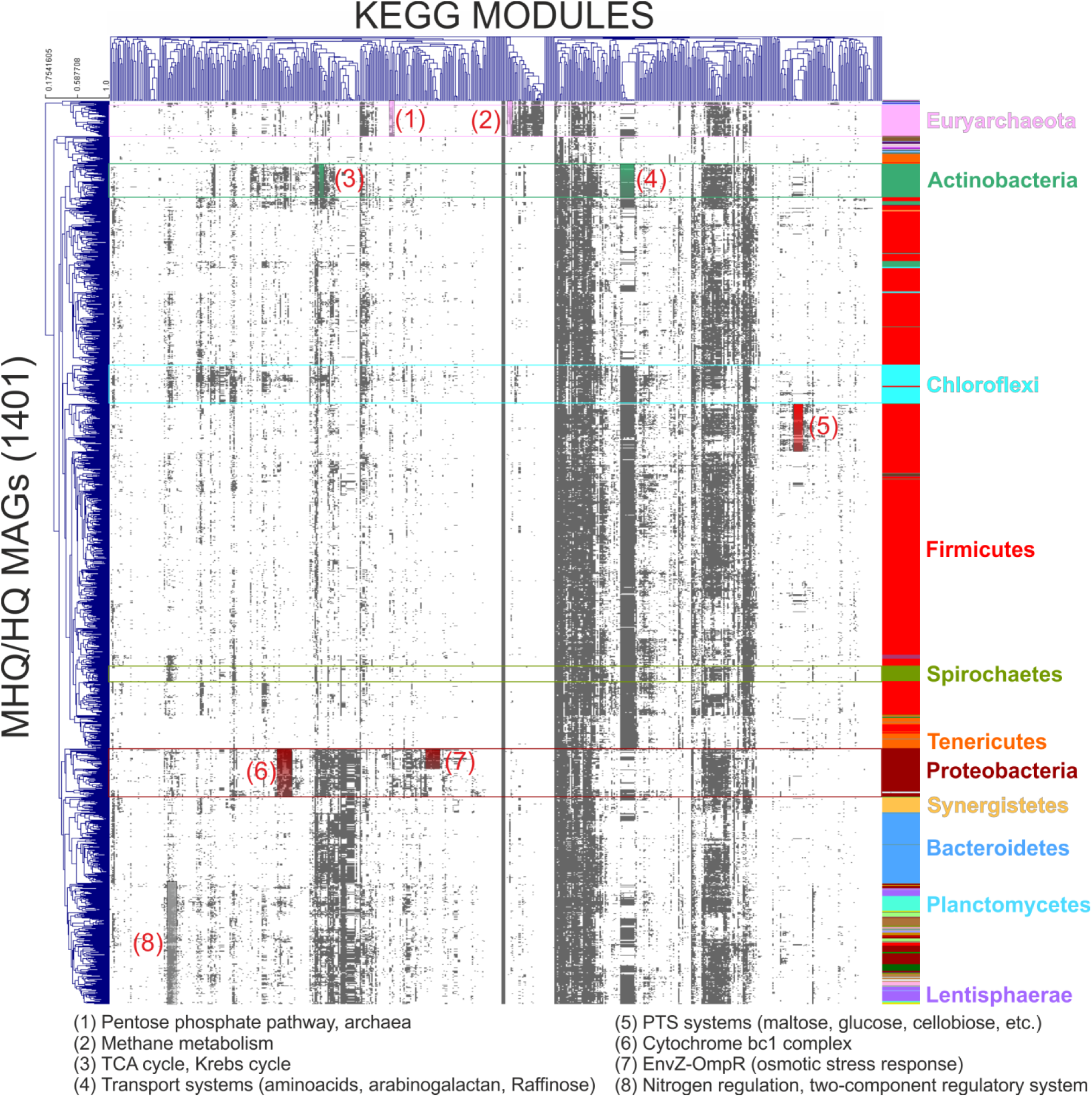
This is a representation of the hierarchical clustering of the “complete” and “1 bm” KEGG modules identified in the HQ and MHQ MAGs (1,401 modules in total). In the right part of the figure MAGs taxonomically assigned at phylum level are shown for the most represented phyla. KEGG modules specifically identified in selected phyla are highlighted and numbered. Colors assigned to the taxa are the same as reported in figure 3.

Initial evaluation was focused on the identification of MAGs having a specific KEGG module. Considering both the complete and “1 bm” modules, only 15 “core modules” have been identified in more than 90% of the HQ-MHQ MAGs. These include for example “C1-unit interconversion”, “PRPP biosynthesis”, “glycolysis, core module involving three-carbon compounds”. Other 223 “soft core modules” were present in 10% to 90% of the HQ-MHQ MAGs. Finally, 289 “shell modules” have been identified in less than 10% of MAGs, including those associated with “methanogenesis”, “reductive citrate cycle” and “Wood Ljungdahl (WL)-pathway”. The high fraction of “soft core” and “shell” modules revealed a highly specialized microbial community, with a small number of species performing crucial functions such as methanogenesis. Considering the “complete” and “1 bm” modules together, the median number of modules per MAG was 107. Results obtained revealed the presence of a small fraction of “multifunctional MAGs” (~1.6%) with more than 180 modules encoded. These microbes are mainly associated to specific taxa, and considering the HQ-MHQ MAGs, they represent 8.6% of the *Proteobacteria*, 14.3% of the *Chloroflexi*, 7.7% of the *Planctomycetes* and only 0.14% of the *Firmicutes*. Thus, the AD microbiome typically comprises “oligofunctional” MAGs, which are characterized by the presence of less than 80 modules. Taxonomic distribution of the 89 HQ “oligofunctional” MAGs demonstrated that they were phyla-specific, representing 91.7% of the HQ *Tenericutes*, 32.2% of the HQ *Euryarcheota* and 19.7% of the HQ *Bacteroidetes*. On the other hand, none of these MAGs was assigned to *Planctomycetes* or *Proteobacteria*. Functional genome classification proposed for MAGs with a low number of modules does not seem to be associated to a limited microbial community distribution, since 14 of these “oligofunctional” MAGs were identified in 70 or more samples (>0.001% abundance). In particular, *Bacteroidales* sp. AS26fmACSIPLY_35 (95.5% Cp) not only is widespread, but frequently represented more than 1% of the microbiome. A higher ability of “multifunctional” MAGs to colonize multiple samples was expected. However, a correlation between the number of modules identified in a MAG and the number of samples colonized was not identified.

Particular attention was given to the modules associated with “methane metabolism”, and especially to the conversion of different substrates (carbon dioxide, acetate, methylamines and methanol) into methane. These modules were identified with different frequencies in the AD microbiome. Carbon dioxide reduction was identified in 29 MAGs (17 encoding complete module), acetate conversion in 25 MAGs (14 encoding complete module), methanol reduction in 40 MAGs (11 encoding complete module) and methylamine-methane conversion in 17 MAGs (12 encoding complete module). The methanogenic archaeon having the highest relative abundance in samples was *Candidatus* Methanoculleus thermohydrogenotrophicum AS20ysBPTH_159 [44]. It was present in 35 samples with abundances higher than 1%, reaching 33% of the total microbiome in a sample where cattle manure was supplemented with lipids (LSBR-R2-with-lipids) (Additional File 7). This MAG encodes complete modules for hydrogenotrophic and acetoclastic methanogenesis. However, additional analysis is needed to verify if this methanogen can use both pathways to produce methane. Other two methanogens frequently found at high relative abundances in the AD system are *Methanosarcina thermophila* AS02xzSISU_89 and *Methanoculleus thermophilus* AS20ysBPTH_14 (2.2% and 1.2% of the microbiome on average). As other members of the *Methanosarcina* genus, AS02xzSISU_89 has a versatile genetic potential to convert different substrates into methane. In accordance, AS02xzSISU_89 encodes all four complete modules of methane metabolism map [45]. In total, 22 methanogens have relative abundances higher than 1% in at least one sample, and 9 higher than 10%.

It is interesting to note that members of the *Thermoplasmatales* class and *Methanomassiliicoccus* order have complete or nearly complete modules for methylotrophic methanogenesis. Functional analysis was only partially possible in *Crenarchaeota, Woesearchaeota* and *Candidatus Lokiarchaeota* due to the difficulty in the identification of the proteins belonging to the methanogenic pathway. However, the low completeness of modules associated to the “carbon dioxide to methane” conversion indicates that these species are most probably restricted to use acetate and/or methylamines.

Apart from the fundamental role of methanogenesis in the AD system, the conversion of acetate, carbon dioxide and hydrogen can follow different pathways and can be strongly influenced by the environmental conditions. Considering the modules associated with carbon fixation, those encountered more frequently were the phosphate acetyltransferase-acetate kinase pathway (acetyl-CoA => acetate) identified in 1,155 MAGs (82.4%) with 988 MAGs encoding the complete module, the reductive acetyl-CoA pathway (also called Wood-Ljungdahl pathway) identified in 86 MAGs (5.8%) with 52 encoding the complete module, and the reductive pentose phosphate cycle (ribulose-5P => glyceraldehyde-3P) identified in 128 MAGs (9.1%) with 42 encoding the complete module. Interestingly, since the WL pathway is present only in 0.49% of the microbial genomes deposited in the KEGG database, in the AD microbiome it is more common than expected. The taxonomic distribution of the 86 MAGs encoding the W-L pathway is mainly restricted to *Firmicutes* (75.6%), followed by *Chloroflexi* (9.3%), *Proteobacteria* (7%), *Euryarchaeota* (3.4%) and *Actinobacteria* (2.3%). Functional activity and syntrophic association with methanogens was previously reported for some of these species (e.g. *Tepidanaerobacter syntrophicus, Syntrophorhabdus aromaticivorans* and *Desulfitobacterium dehalogenans)* [46–48]. However, the vast majority was not previously characterized at the genome level, suggesting that potential syntrophic associations or acetogenic behavior are present in many unknown species. Most of the MAGs encoding the W-L pathway are rare in the microbiome and on average they do not exceed 1% of relative abundance. However, under certain conditions they can become dominant, as for example *Firmicutes* sp. AS4GglBPBL_6 (24.8% relative abundance in the Fangel biogas plant), *Firmicutes* sp. AS02xzSISU_21 (32% in reactor fed with avicel) and *Firmicutes* sp. AS4KglBPMA_3 (12% in the Nysted biogas plant). Globally, 17 of the MAGs with W-L pathway exceeded 1% abundance in the AD microbiome in at least one sample. Interestingly, the Fangel biogas plant showed a high total ammonia level during the sampling process (4.2 g/L) (Additional File 1). This indicates that, despite SAO bacteria are usually present at low abundances, environmental parameters of the reactors can strongly influence their abundance and probably their activity, as also previously suggested [8]. Despite it is hard to classify these species as SAO or acetogens. This result can provide a more accurate evaluation of the fraction of bacteria involved in acetate conversion and can support the delineation of a more accurate mathematical model for the AD process.

Considering the relative percentage of HQ MAGs in each condition, along with the completeness of KEGG modules, it was possible to estimate the relative abundance of each module in all samples (Additional File 10). Although measurements at the RNA/protein level are needed to have direct information on pathways activity, it is evident that different samples have highly variable representation of crucial KEGG modules (Fig. 6). It is interesting to note that the relative abundance of MAGs potentially associated to the hydrogenotrophic and acetoclastic methanogenesis is highly variable among samples. Particularly, in biogas plants characterized by low TAN (1.9-2 mg/L) (e.g. “BP-Gimenells” and “BP-LaLlagosta”), acetoclastic methanogenesis is favored and the ratio acetoclastic/hydrogenotrophic is 0.94 and 0.99, while in biogas plants where TAN is high (4-7 mg/L) (e.g. “BP-Vilasana”, “BP-Torregrossa” and “BP-Fangel”) the ratio acetoclastic/hydrogenotrophic is 0.16, 0.21, 0.02. Analyzing reactors where ammonia levels were reported, it was found indeed a significant correlation (R^2^ 0.62, p 9.3 E^−5^) between ammonia concentration and the “acetoclastic/hydrogenotrophic” ratio. Additionally, there is a high level of acetoclastic methanogenesis in reactors fed exclusively with acetate, such as “LSBR-D122-DNA-BF-Rep1”, “LSBR-D200-DNA-BF-Rep1” and “LSBR-R3-acetate”. Relative abundance of the methanogenic modules was found to be highly different among samples considered. As expected, it was close to zero in acidogenic reactors (pH<5, “LSBR-DSAc-preH_2_” and “LSBR-DSAc-postH_2_”) and very high in reactors with acetate as feeding substrate (e.g. “LSBR-D200-DNA-BF” or “LSBR-R1-acetate”). The high abundance of methanogenic modules in the latter reactors can be correlated with the direct use of the substrate by acetoclastic methanogens, with a parallel reduction of the species encoding the W-L pathway.

**Fig. 6.**
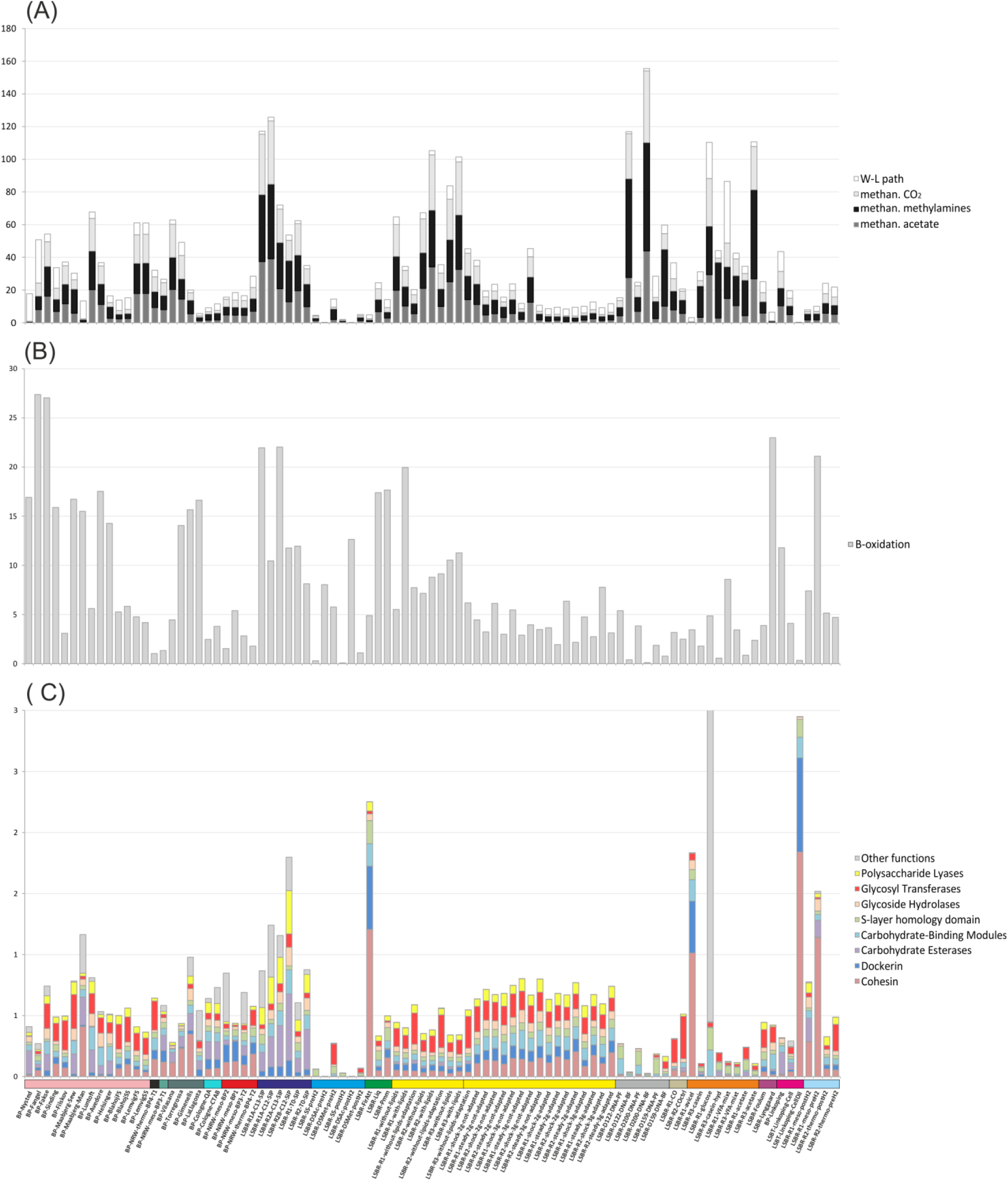
Representation of the relative abundance of four relevant functional modules in the AD system: “methanogenesis from CO_2_”, acetate and methylamines, “W-L pathway”. Bar graph was obtained for each sample by summing the relative abundance of all the HQ and MHQ MAGs encoding these “complete” or “1 bm” modules. In figure samples collected from biogas plants were reported in the left part of the figure (first 26 samples), while those derived from laboratory reactors or batch tests were shown in the right part of the figure.

Cellulosic biomass in AD is represented by agricultural residues and dedicated energy crops, and is the most abundant carbon source [49]. Previous findings suggested that many microbes participating to the AD food chain are involved in hydrolysis of polysaccharides, which is a crucial step in organic matter degradation [30]. In order to find the species involved in complex carbohydrate decomposition, MAGs featuring high enrichments in CAZymes (p<1*e-5) have been selected for further analysis (Additional File 11). A specific MAG was frequently found to be enriched in more than one functional class, e.g. 35 MAGs are enriched in “carbohydrate esterases”, 32 of them were also enriched in “glycoside hydrolases” and 13 in “polysaccharide lyases”. Eleven MAGs are enriched both in enzymes having cohesin and dockerin domains, and additionally other 21 have high enrichment in only one of these two classes. A selective enrichment in one class (dockerins or cohesins) is quite common in microbial species [50]. Globally, 490 HQ MAGs (35% of the total) are enriched in one or more CAZymes classes, evidencing that polysaccharide degradation is one of the most widespread functional activities in the AD system. Although polysaccharide degraders are frequently associated to *Firmicutes* (246 MAGs) and *Bacteroidetes* (68 MAGs), many other phyla were found to be enriched and an involvement in polysaccharide degradation can be hypothesized for members of other taxa. For example, all MAGs belonging to the Candidatus *Hydrogenedentes*, the *Armatimonadetes*, 90% of the *Fibrobacteres*, 93% of the *Lentisphaerae* and 85% of the *Planctomycetes* are potentially involved in this process. Some members of the CPR taxa are also predicted as associated to carbohydrate degradation, such as Candidatus *Dojkabacteria* and Candidatus *Pacebacteria*.

MAGs enriched in specific classes of CAZymes have a higher relative abundance in specific samples (Fig. 6 C); as expected, MAGs encoding for cellulosomes, are frequent in reactors fed with polysaccharides (avicel or cellulose, LSBR-R1-avicel and LSBT-Linkoping-CellDil) or in samples enriched with microbes firmly attached to the plant material, e.g. LSBR-FirAtt [30,51]. The cellulose-degrading MAGs have a scattered distribution among these samples; for example *Clostridium sp. AS05jafATM_73* has an impressive relative abundance of 23.8% in LSBT-Linkoping-CellDil but is very rare in LSBR-R1-avicel, where it is replaced by strictly related species, e.g. *Ruminiclostridium thermocellum AS02xzSISU_19* and *Clostridium* sp. *AS20ysBPTH_178*. These findings suggest that degradation of polysaccharides is one of the main driving forces expanding the complexity of the AD microbiome, since polysaccharide degradation can be alternatively performed by many different microbes having a scattered distribution in different samples.

A tentative estimation of the relative impact of the polysaccharide degradation process in different samples (Fig. 6 C) was obtained by considering the relative abundance of MAGs encoding genes for a specific function (e.g. “cohesin”, “dockerin”, or “Carbohydrate Esterases”). A few samples are dominated by polysaccharide hydrolyzing MAGs, (e.g. “LSBR-R1-avicel”), most probably because they were fed with substrates rich in cellulose, while generally the fraction is lower than 2%, particularly in biogas plants. This indicates that, despite the number of MAGs involved in polysaccharide degradation is high, the relative abundance of most species is low. This can be due to the presence of relative minor players in terms of abundance, but having high transcriptional activity; if they are highly active, they can enhance or trigger the metabolic processes of dominant members. However, this needs additional verification to be demonstrated. Rectors fed with blended manure (LSBR-R2B-C12-SIP) and with glucose only (“LSBR-R1-glucose”) have a high fraction of species with genes encoding carbohydrate esterases and glycosyl transferases. On the other hand, reactors fed with simplified substrates such as acetate (“LSBR-D122-DNA-BF” and “LSBR-D200-DNA-BF”) or cheese whey (“LSBR-DSAc-preH2”, “LSBR-SS-postH2”) have a negligible fraction of polysaccharide-degrading species.

### MAGs replication rate

To our knowledge, there are no known studies in the literature reporting on an extensive evaluation of growth dynamics for microbial species of the AD system. To determine the replication rate of MAGs across multiple samples, the sequencing coverage resulting from bi-directional genome replication was used to calculate the index of replication (iRep) [52]. In total, 2,741 measurements (1986 after calculation of average values over replicates) were obtained for 538 MAGs (Additional File 12). Considering the average iRep values determined in all different samples for each MAG, it was obvious that nearly 90% of species showed similar values between 1.1 and 2, and only 10% had values between 2 and ~4 and can be considered as “fast growing”. Among them are microbial members of the poorly characterized phylum *Atribacteria* (Candidatus *Atribacteria* sp. AS08sgBPME_53, iRep 2.9), the thermophilic protein degrader *Coprothermobacter proteolyticus* AS01afH2WH_35 (iRep 2.8) and the candidate syntrophic species *Defluviitoga tunisiensis* AS05jafATM_34 (iRep 2.6) [53]. iRep values for 301 MAGs were obtained for various conditions, allowing an evaluation of possible influences of environmental parameters on their replication rates. Results were obtained for 28 phyla evidencing that *Tenericutes, Spirochaetes*, Candidatus *Atribacteria, Thermotogae, Synergistetes*, and *Coprothermobacterota* have on average high iRep values (iRep 1.81, 1.85, 2.12, 2.27, 2.42, 2.84, respectively), while *Euryarchaeota* and *Acidobacteria* have low values (1.45 and 1.46) (Fig. 7 A). MAGs belonging to phyla *Bacteroidetes* and *Firmicutes* have similar (and low) iRep values (1.69 and 1.53 on average) except some outliers. Otherwise, iRep values assigned to *Synergistetes* and *Coprothermobacterota* are distributed over a wide range, but on average are higher than for other phyla (2.4 and 2.8) (Fig. 7). The limited growth rate of some taxa, such as *Acidobacteria*, was also previously reported [54] and it was speculated that this property hampered their isolation. The high iRep values measured here for some known species also suggests that their isolation may be easier as previously assumed [55]. The low replication rate of *Euryarchaeota* (~1.45 on average) is also known [56], but in the present study, comparative data for 17 MAGs were reported extending previous findings and providing results for multiple species having different abilities in substrates utilization. In particular, the *Euryarchaeota* with the highest iRep is *Methanothrix soehngenii* AS27yjCOA_157, followed by *Methanomicrobiales* sp. AS21ysBPME_11 (Fig. 7 B). *M. soehngenii* was previously defined as a slow-growing methanogen specialized to utilize acetate [57] and it is very interesting that 7 out of 9 iRep results obtained are higher than 2, while the highest value obtained for *Methanosarcina thermophila* was 1.7. The replication rate of AS21ysBPME_11 was determined in a limited number of samples associated with key long-chain fatty acid-degrading populations (e.g. “LSBR-R1A-C13-SIP”), while in other samples associated with UASB reactors, it had a low replication rate. Findings reported for AS21ysBPME_11 indicate that, in a complex microbiome, growth rates can be very different compared to those obtained for isolated species under laboratory conditions, possibly because of cooperative/syntrophic associations with other microbes, or difficulties in identifying the appropriate growth medium.

**Fig. 7.**
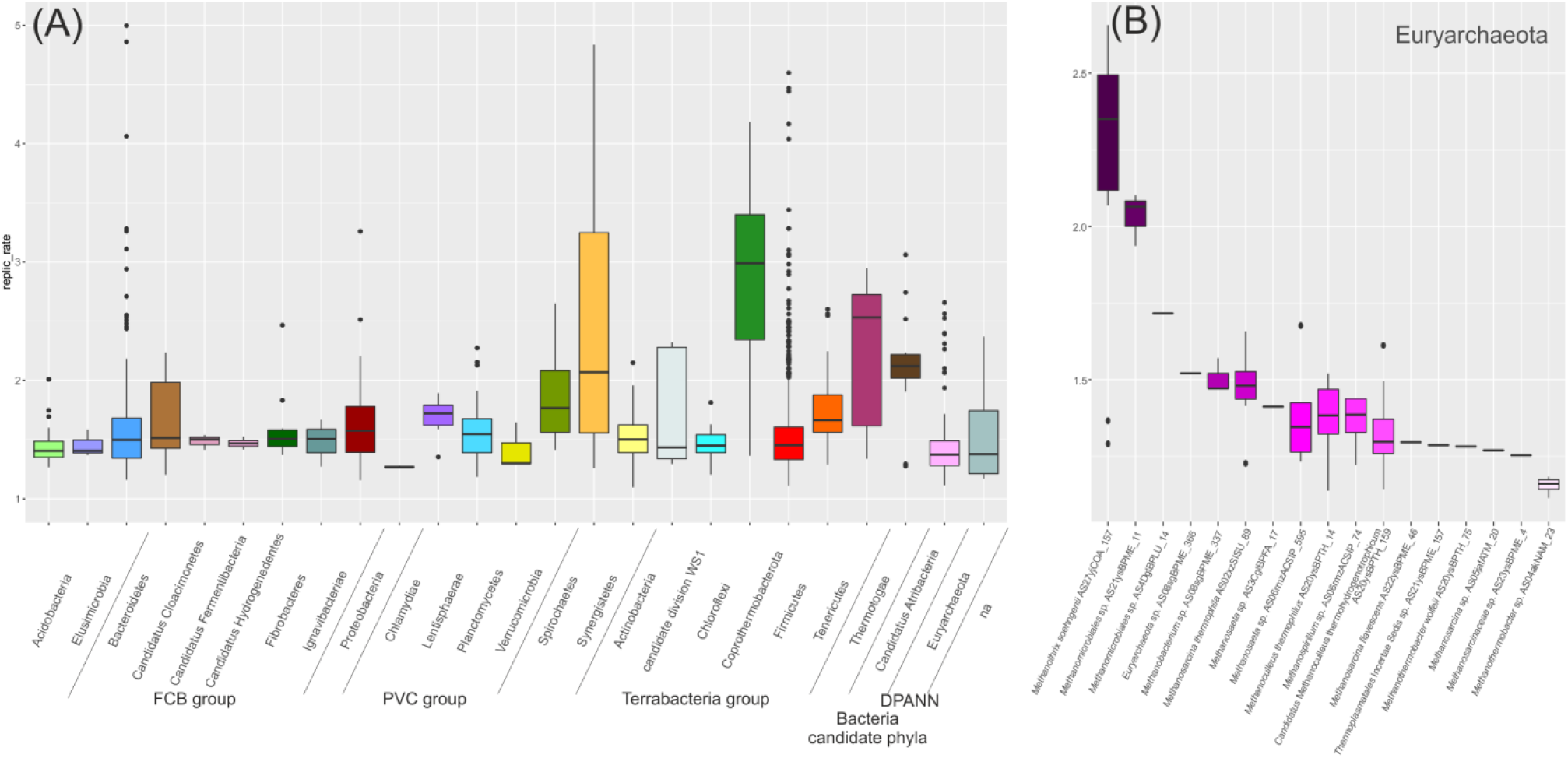
(A) Distribution of iRep values obtained for 538 MAGs are reported considering separately MAGs belonging to each of the 25 phyla having at least three MAGs (“na” refers to taxonomically unassigned MAGs). (B) Distribution of iRep values obtained for Euryarchaeota. MAGs having only one value are reported as a horizontal bar.

Our findings also suggest that duplication rates are dependent on metabolic properties of MAGs. Calculation of iRep values performed independently for MAGs encoding different KEGG modules evidenced that MAGs involved in polysaccharide degradation have quite low iRep values; this is more evident for microbes growing attached to plant material with cohesin/dockerin domains (iRep 1.43). These species represent the so-called slow-growing cellulolytic microflora [58], and include for example *Clostridium* sp. AS05jafATM_73 and *Clostridium* sp. AS24abBPME_43 (iRep 1.2 and 1.4). Those having “carbohydrate-binding modules” (iRep 1.48) and “glycosyl transferases” (iRep 1.48) have also a slow growth rate. Species involved in “energy metabolism - methane metabolism” have low iRep values (1.46, 1.45 and 1.45), while those associated to “carbon fixation” (e.g. “reductive citrate cycle” or “W-L pathway”) have higher values (iRep 1.55; 1.56). MAGs encoding the “reductive pentose phosphate cycle” have even higher values (iRep 1.72). Although generally low, iRep values were also obtained for poorly characterized taxa such as Candidatus *Atribacteria* and Candidatus *Fermentibacteria* (Fig. 7 B), suggesting that they are slow-growing members of the AD system.

Availability of iRep values for a large number of species, and their association with functional roles of microbes can provide an estimate of the growth rate of species involved in particular steps of the AD food chain. Since nowadays mathematical models of the AD system are based on growth rates measured for a limited number of species, information obtained from iRep can provide a more generalized representation of microbial dynamics which can be included in simulations, reinforcing their predictive efficiency.

## Conclusions

A comprehensive genome-centric assessment of the AD microbiome was obtained here starting from 134 samples downloaded from public databases. Results obtained at the species-level suggest that the biogas microbiome is extremely flexible, allowing it to adapt to the different conditions present in different biogas reactors, including a wide range of temperatures and substrates. This adaptation is facilitated by the presence of multiple different microbial communities that have little to no overlap among them. Considering the abundant MAGs, only 25 were identified in numerous samples. However, there are many other MAGs which are present at low abundances constituting a persistent but low-abundance microbiome. Our findings related to metabolic pathways showed a partitioning of microorganisms according to their predicted substrate utilization capacities.

Investigation of metabolic pathways suggested that some crucial processes, such as conversion of acetate to CO_2_, may be performed by a limited number of species. The high heterogeneity regarding protocols used for sample collection, DNA extraction, sequencing and metadata registration evidenced that common protocols are direly needed to obtain datasets that can more easily be compared among different experiments and working groups. By reconciling numerous metagenomics studies previously reported in the literature, this study suggests that the establishment of a global database of genomes is of great importance for future studies.

## Methods

### Selection of samples and reads filtering

Illumina sequences were downloaded from Sequence Read Archive (SRA), MG-RAST or JGI Genome portal databases. Quality check and adaptors removal were performed using Trimmomatic (v0.33) (LEADING:20 TRAILING:20 SLIDINGWINDOW:4:20 MINLEN:70) [59] and bbduk (version released Nov 2016) (https://jgi.doe.gov/data-and-tools/bbtools/). In total, 134 samples have been selected, which was reduced to 91 after merging biological replicates. The composition of the feedstocks used in the different reactors was approximated using substrate information from various sources (Additional File 1). When available, such data was taken from the publicly accessible description of the respective experiments or full-scale plant operation datasets. Otherwise, reactor feedstocks were estimated by a proportionality-based mixing technique: taking the characteristics of their individual constituents from available literature and combining them according to their fresh matter (FM) or volatile solid (VS) ratio in the feed.

### Assembly

Reads obtained from biogas reactors inoculated with the same inoculum were co-assembled, as well as those collected from primary and secondary reactors of the same biogas plant. On contrary, samples derived from reactors using different inocula and those collected once from a specific biogas plant were assembled individually. Before performing the co-assembly of reads obtained from different reactors, the similarity of the microbial composition among different samples was verified running MetaPhlAn2 (v2.2.0) on one million unassembled reads, randomly collected from each sample [60]. This preliminary check confirmed that samples collected from the same reactor, or collected from reactors using the same inoculum had on average similar microbial composition. Reads were assembled using Megahit (v1.1.1) with “--sensitive” mode for samples having less than 40 Gb of sequenced bases and with “--large” for the remaining assemblies [61]. Quality of the assemblies was determined using QUAST (v3.1) [62] and results are reported in Additional File 2.

### Binning

Applying the Bowtie 2 program (v2.2.4) [63] a trial alignment was performed using 100,000 randomly selected reads per each sample in order to calculate the fraction of reads aligned on each assembly. This allowed the identification of all samples having a reasonable alignment rate on each assembly (higher than 25%) and to select them for the subsequent binning step. Samples having less than 25% aligned reads were considered as being not informative. Based on these preliminary results, the number of experiments considered for coverage calculation and subsequent binning ranged from 11 to 89 depending on assembly. Using MetaBAT 2 (v2.12.1) [64] bam files were inspected and each assembly was binned using standard parameters. Minimum size of scaffolds considered for MAGs generation was 1.5 kbp. MAGs were checked for completeness (Cp) and contamination (Ct) using the “Lineage_wf” workflow of CheckM (v1.0.3) [65] and the result obtained for each MAG was determined using the formula: CC3=Cp-(Ct*3). Removal of contamination from MAGs was performed using RefineM (v0.0.23) [37]; contaminating scaffolds were identified considering their genomic characteristics (GC content and tetranucleotide composition). After the filtering step performed with RefineM, the “CC3 value” of each MAG was calculated again leading to only 159 MAGs showing an improved “CC3 value” after contamination removal; all the remaining MAGs were maintained in their initial condition (without performing the filtering step).

### MAGs de-replication

MAGs obtained were de-replicated using a double step process based on a fast Mash software (v2.0) verification [66] performed on the entire genome sequences and using very permissive parameters (0.05 Mash-distance, roughly equivalent to 0.95 ANI and 100/1000 Matching-hashes). Subsequently, a more precise analysis was performed applying the genome-wide Average Nucleotide Identity metric (ANI) [67] using protein-encoding nucleotide sequences only. ANI calculation was performed only on the comparison satisfying the first Mash verification step. MAGs were considered as belonging to the same species if they were having ANI higher than 95% and reaching at least 50% of genome coverage for both strains (Varghese *et al*., 2015) (on at least one of the two comparisons, “MAG1 vs. MAG2” or “MAG2 vs. MAG1”). After ANI calculation, from each cluster of MAGs which belong to the same species, a representative one with the highest CC3 value was selected. These MAGs were classified in three groups according to their quality and contamination levels: High Quality “HQ” (Cp>90%, Ct<5%), Medium-High Quality “MHQ” (90%>Cp>=70%; 5%<Ct<10%) and Medium Quality “MQ” (70%>Cp>=50%; 5%<Ct<10%).

### Taxonomic assignment

Taxonomic classification was determined for 1,635 MAGs obtained after de-replication and belonging at least to the MQ level. This approach was carried out as described previously [7] with small modifications: (1) The highest priority for taxonomy assignment has been given to the ANI results obtained comparing MAGs with genomes from NCBI database. gANI calculation was performed as described below comparing MAGs and the genomes downloaded from NCBI microbial genome database (last accessed date: May, 2018). 56 MAGs showed an ANI value higher than 95% and more than 70% of genes in common with the reference species. Other 149 MAGs were also highly similar to known species deposited at the NCBI microbial genome database, but these reference genomes were not taxonomically assigned at species level. Other 38 MAGs had average similarity which was higher than 95%, but the percentage of common genes ranged between 50% and 70%. Furthermore, affiliation of these microbes to the genus level was doubtful. (2) Intermediate priority for taxonomy classification was given to MAGs encoding the 16S rRNA genes longer than 300 bp. The 16S rRNA genes were identified for each MAG with in-house developed perl script using Hidden Markov Models obtained from RNAmmer [68] and taxonomy assessment was determined using RDP classifier trained on SILVA 132 ribosomal RNA (rRNA) database [69]. Taxonomy results were compared with those obtained from ANI and from taxonomically informative proteins (PhyloPhlAn and CheckM, “step 3” below). Five discordant results were manually verified and corrected removing possibly misassigned 16S rRNA genes. (3) Results obtained from taxonomically informative proteins (PhyloPhlAn and CheckM) were used for taxonomic classification of the remaining MAGs. Finally, results obtained applying all three methods were compared with each other in order to discover discrepancies, which were identified and manually corrected only for the MAG *Candidatus* Fermentibacter daniensis_AS4DglBPLU_32. An additional verification was performed on MAGs assigned to CPR, DPANN and some other hypothetical taxa by selecting 5278 representative genomes from NCBI microbial genomes database as described previously [7], building a tree using PhyloPhlAn [70] and performing a manual inspection assisted by Dendroscope (v1.4) [71]. MAGs were classified against all taxonomically classified taxa of the NCBI Genome Database (prokaryotic section) using Microbial Genomes Atlas MiGA Online [41]. Results obtained were compared with those determined using the combination of methods describe at the beginning of this section.

### MAGs coverage calculation and relative abundance

Filtered shotgun reads randomly selected from each sample were aligned back to the entire collection of MAGs. Ordered “bam” files were inspected using CheckM [65] in order to calculate both, the fraction of reads aligned on the recovered genomes and the relative abundance of each MAG in all the samples. Analysis was performed using all reads available for each sample and verified using a representative subsample of one million reads per sample. Results obtained using the two datasets of sequences were highly similar (correlation coefficient was >0.999 on MAGs representing more than 0.001% of the population). Results obtained using one Mreads per sample are reported in Additional File 7. The value (0.001%) was also defined as the arbitrary threshold for considering one MAG as “present in a specific sample”. Coverage values obtained for each MAG were clustered with MeV (v4.9.0) using Pearson correlation and average linkage [72]. The fraction of MAGs shared between different samples was visually represented using CIRCOS (v0.69) [73]. Alpha and beta diversity were determined from the file reporting the number of reads per MAG on each experiment applying Past (v3.21) [74]. The same tool was used for statistical tests and graphical plots.

### Gene finding and annotation

Gene annotation was performed using three different procedures: (1) Protein-encoding genes were predicted and annotated using Rapid Annotation Using Subsystem Technology (RAST annotation server) [75]. This SEED-based annotation was also used to predict rRNA genes and tRNAs. The number of protein encoding genes belonging to each subsystem was determined as previously described [7]. These results were reported in a table for comparative purposes (Additional File 13). (2) Protein-encoding genes were predicted using Prodigal (v2.6.2) run in normal mode and associated to KEGG IDs using Diamond (v0.9.22.123) [76]. KEGG IDs were associated to modules to determine completeness using “KEGG Mapping/Reconstructmodule.py” (https://github.com/pseudonymcp/keggmapping). Abundance of all the KEGG modules in each experiment was calculated with custom perl scripts considering pathway completeness and relative abundance estimated for all MAGs (https://sourceforge.net/projects/perl-scripts-kegg/). Cluster analysis on “complete” or “1 bm” KEGG modules identified in HQ and MHQ MAGs was performed using MeV (v4.9.0) [72]. (3) To predict genes encoding carbohydrate-active enzymes, the carbohydrate-active enzyme database (CAZy) annotation web server dbCAN [77] was used taking as input proteins predicted with Prodigal (v2.6.2) run in normal mode. In total 305 classes of carbohydrate-active enzymes were identified, which were clustered in ten groups (Additional File 11). Abundance of specific functional classes was determined using hypergeometric analysis and p-values corrected using false discovery rate as described previously [78].

### MAGs replication rate

Considering the genome size and the total number of reads mapped on each MAG, the coverage of each MAG was determined using Bowtie 2 (v2.2.4). The MAGs having completeness higher than 90%, contamination lower than 5%, a number of scaffolds per Mbp lower than 175 and a coverage value higher than five, were selected in order to determine their index of replication (iRep) applying the iRep software [52].

### β-diversity and statistics

β-diversity was calculated applying the ExpressBetaDiversity (EBD) software (v1.0.7) [79]. Statistical calculations, α and β-diversity calculation were performed using past software (v3.21) [74].

## List of abbreviations

(MAG): Metagenome Assembled Genome
(AD): Anaerobic Digestion
(Cp): Completeness
(Ct): Contamination
(ANI): Average Nucleotide Identity
(HQ): High Quality
(MHQ): Medium-High Quality
(PCoA): Principal Coordinate Analysis
(WL): Wood–Ljungdahl
(iRep): Index of Replication
(FM): Fresh Matter
(VS): Volatile Solids

## Declarations

### Ethics approval and consent to participate

(Not applicable)

### Consent for publication

(Not applicable)

### Availability of data and material

Shotgun sequences used were downloaded from SRA, EBI, DDBJ, GJI or MG-RAST and all the information associated to the projects are reported in (Additional File 1). All the MAGs sequences are available through the MiGA database under the project “http://microbial-genomes.org/projects/biogasmicrobiome” and in the biogasmicrobiome website (https://biogasmicrobiome.env.dtu.dk/).

### Competing interests

The authors declare that they have no competing interests.

### Funding

This work was supported by the Innovation Fond under the project “SYMBIO—Integration of biomass and wind power for biogas enhancement and upgrading via hydrogen assisted anaerobic digestion,” contract 12-132654.

### Authors’ contributions

S.C., I.A., L.T., designed experiments. S.C., L.T., X.Z., A.K., L.R.R., R.M.Z., A.B. and K.T.K. analyzed the data. S.C., L.T., I.A., I.M., A.S., P.G.K., R.Z. and G.L. wrote the manuscript.

## Acknowledgements

We would like to acknowledge all the people that generated and submitted the sequencing data to public databases, and that were involved in all the projects considered here for the reconstruction of the global AD microbiome.

## Additional files

Additional figure S1: Relative abundance of all the MAGs associated to the phyla identified is represented as a heatmap (see color scale at the bottom of the figure). On top of the figure color codes are associated to the different experiments considered. From top to bottom, phyla are ordered considering their average relative abundance in all the experiments.

Additional figure S2: Cluster analysis of the MAGs relative abundance values. White “cells” represent undetected MAGs, the remaining cells are reported according to a color scale with values increasing from blue to red (see color scale at the bottom). In the right part of the figure colors refer to the taxonomic assignment of the MAG at phylum level. Clusters of MAGs (G1, G2) and clusters of experiments (C1-C3) are discussed in the text.

Additional figure S3: Principal Coordinate Analysis (PCoA) performed considering MAGs abundance in the samples examined. Samples are colored according to the temperature (A), to the feedstock (B) and to the experiment (C). Feedstock composition was summarized according to the data reported in Supplementary data 1. Full-scale biogas reactors are reported as small squares, while laboratory-scale reactors as circles.

Additional figure S4: (A) Dots represent MAGs relative abundance in the samples examined. Dots are colored according to the taxonomic assignment of the MAGs at phylum level. (B) Number of samples where each MAG was identified at relative abundance higher than 0.001%. Each dot is representative of a single MAG and the number of samples where it was identified is reported in y axes (CPR are marked with asterisks).

Additional file 1: Metadata of the samples included in the present study. The composition of the feedstocks used in the different reactors was approximated using substrate information from various sources. When available, such data was taken from the publicly accessible description of the respective experiments or full-scale plant operation datasets. Otherwise, reactor feedstocks were estimated by a proportionality-based mixing technique: taking the characteristics of their individual constituents from available literature and combining them according to their fresh matter (FM) or volatile solid (VS) ratio in the feed.

Additional file 2: Global statistics of the assemblies and binning.

Additional file 3: MAGs taxonomy. MAGs obtained after redundancy removal (based on ANI calculation) were assigned to the taxonomy using three different methods and results were subsequently combined.

Additional file 4: Comparison of MAGs with those reported in previous projects.

Additional file 5: Relative abundance of each taxa in the samples examined. Relative abundance of all the MAG having the same toxonomic assignment were combined in order to determine the relative abundance of each taxon as reported in columns D-CP. All the txonomic levels from kingdom to genus were considered.

Additional file 6: Diversity indexes. Indexes were calculated for each sampl in order to estimate the characteristics of the microbiome. Results were obtained starting from the files reporting the number of reads assigned to each MAG on each sample (subsampling all the samples to 1 million reads) and subsequently elaborated using PAST software. Colors reported in line 2 refer to the projects from which reads have been collected. Remaining colors were assigned in order to discriminate low and high values in a heatmap scale.

Additional file 7: MAGs coverage per sample. MAGs coverage was calculated on all the samples (average values were considered for samples collected in replicates).

Additional file 8: KEGG modules. Completeness of each KEGG module was reported for the high and medium-high quality MAGs.

Additional file 9: MAGs having KEGG modules. Number of MAGs having each KEGG module are reported. Only MAGs having complete and 1 block missing KEGG modules are considered. In column B is reported the total number of MAGs per each phylum (considering only high quality and medium-high quality MAGs). KEGG modules are reported in columns.

Additional file 10: KEGG pathways abundance in different microbial samples. Abundance of all HQ and MHQ MAGs having a complete or “one block missing” KEGG module were summed and the value is reported here as an esitmate of the abundance of this pathway in each sample.

Additional file 11: Enrichment of CAZymes classes in MAGs. Number of proteins assigned to different classes and determined using dbCAN software. Only HQ and MHQ MAGs have been considered.

Additional file 12: Index of replication. iRep values were calculated for each MAG in all the samples considered. Average values were calculated for replicate samples. All samples are reported in column 2, while MAGs IDs and names are reported in columns A and B.

Additional file 13: Resuls obtained using RAST (Rapid Annotation using Subsystem Technology). MAGs genes were assigned to the function using RAST and the number of genes identified per each functional category (FL=firstlevel; SL=second level) are reported. Functional categories are reported in column B, while MAGs are reported in rows (2 is MAG ID and 3 is MAG name assigned considering taxonomy).

## Notes

http://microbial-genomes.org/projects/biogasmicrobiome

https://biogasmicrobiome.env.dtu.dk/

