## Additional Fig for "The anaerobic digestion microbiome: a collection of 1600 metagenome-assembled genomes shows high species diversity related to methane production"

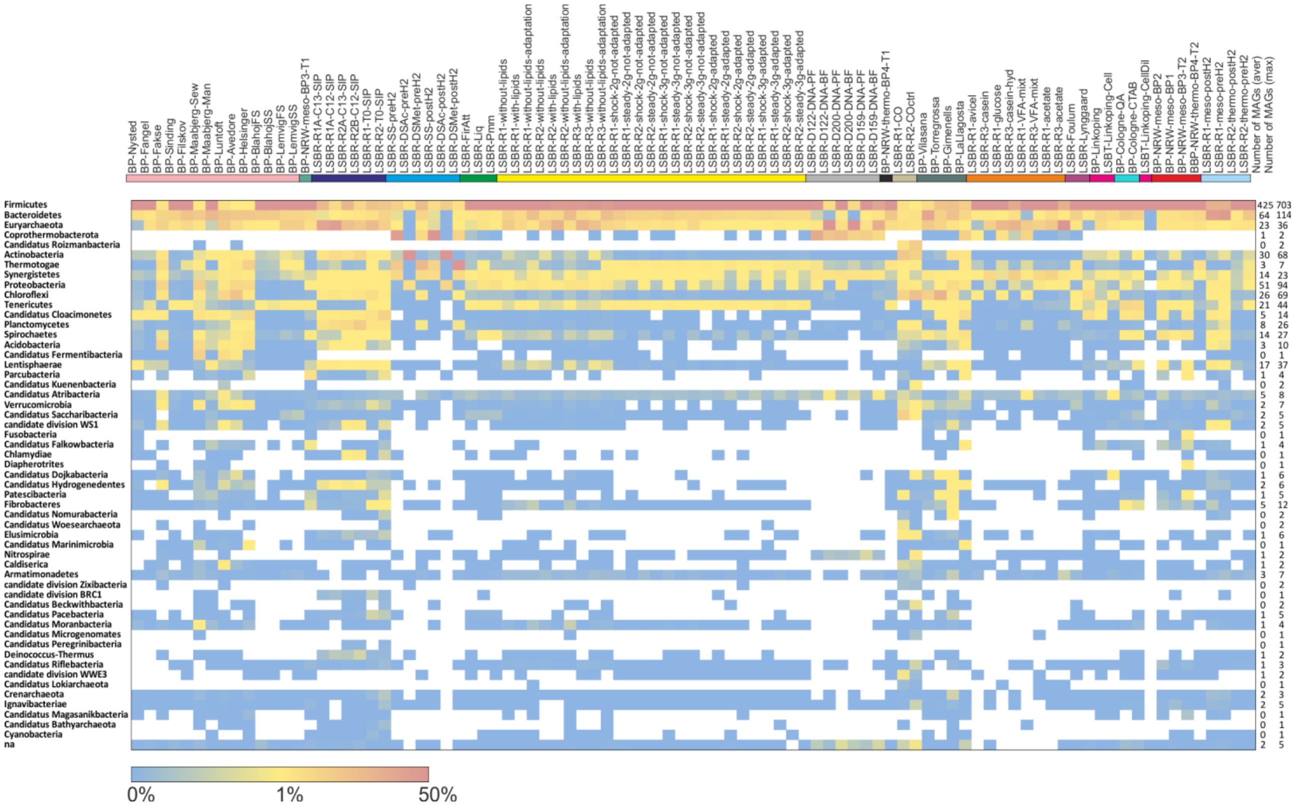


**Additional Fig. S1.** Relative abundance of all the MAGs associated to the phyla identified is represented as a heatmap (see color scale at the bottom of the figure). On top of the figure color codes are associated to the different experiments considered. From top to bottom, phyla are ordered considering their average relative abundance in all the experiments.

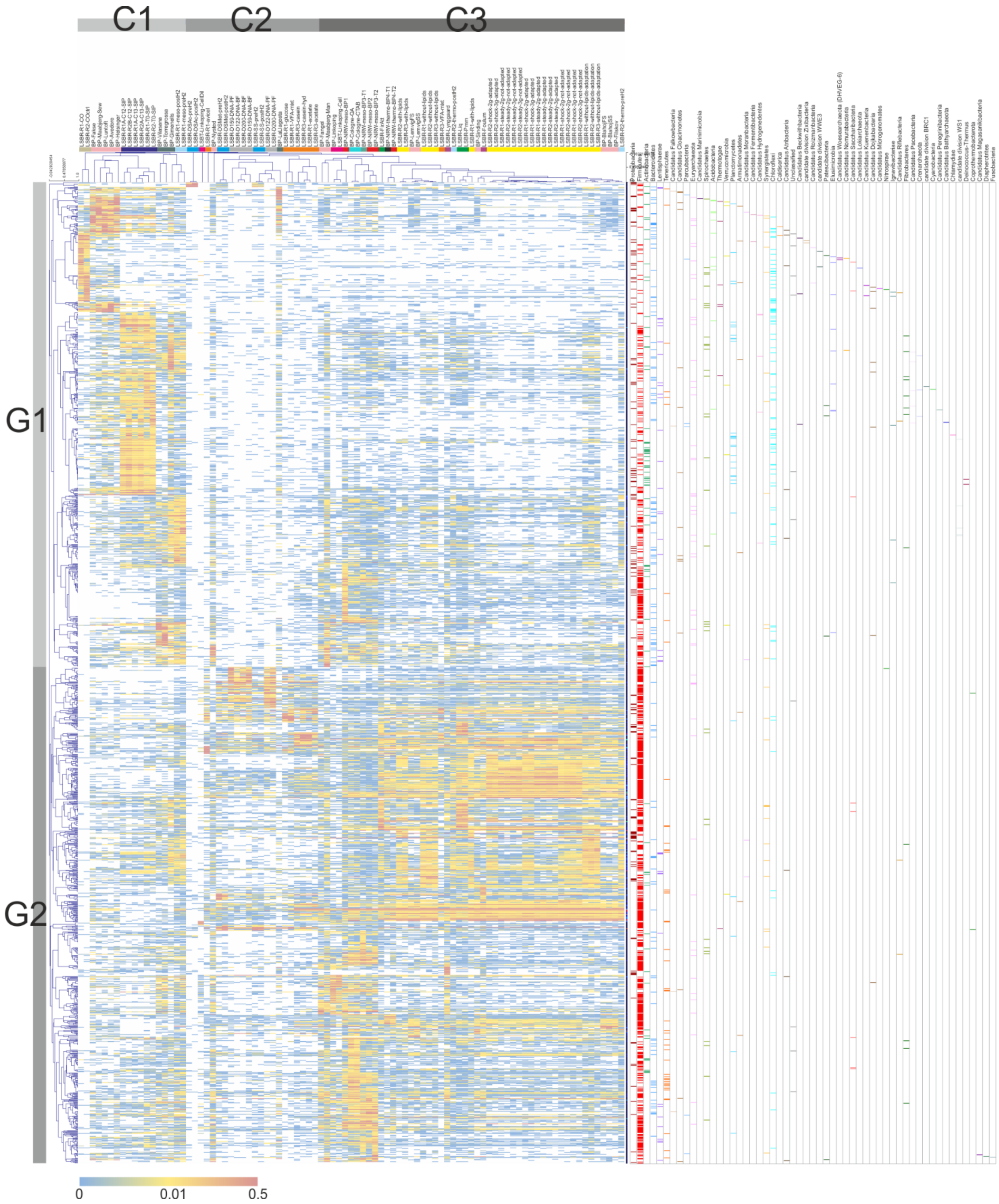


**Additional Fig. S2.** Cluster analysis of the MAGs relative abundance values. White "cells" represent undetected MAGs, the remaining cells are reported according to a color scale with values increasing from blue to red (see color scale at the bottom). In the right part of the figure colors refer to the taxonomic assignment of the MAG at phylum level. Clusters of MAGs (G1, G2) and clusters of experiments (C1-C3) are discussed in the text.

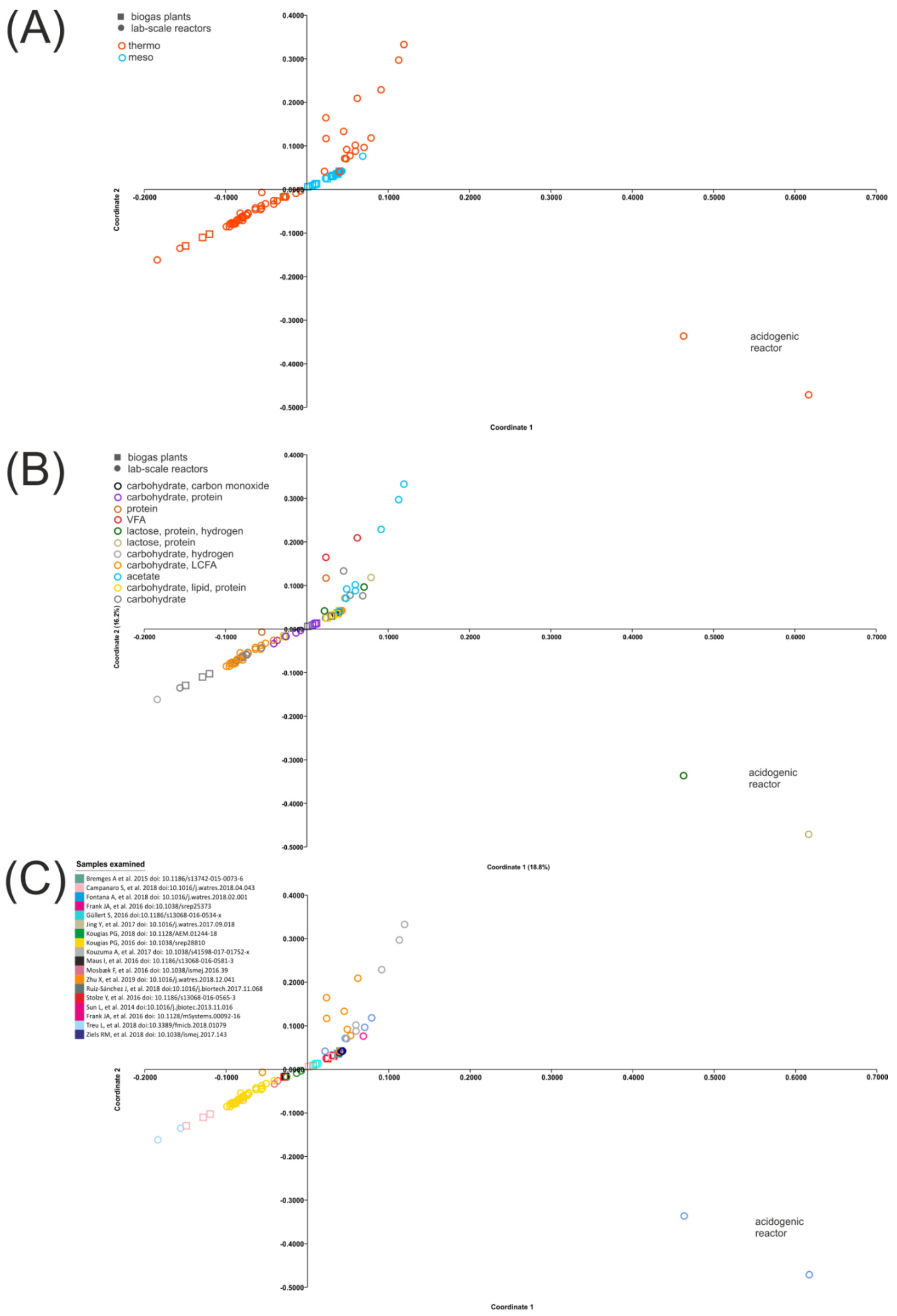


**Additional Fig. S3.** Principal Coordinate Analysis (PCoA) performed considering MAGs abundance in the samples examined. Samples are colored according to the temperature (A), to the feedstock (B) and to the experiment (C). Feedstock composition was summarized according to the data reported in Supplementary data 1. Full-scale biogas reactors are reported as small squares, while laboratory-scale reactors as circles.

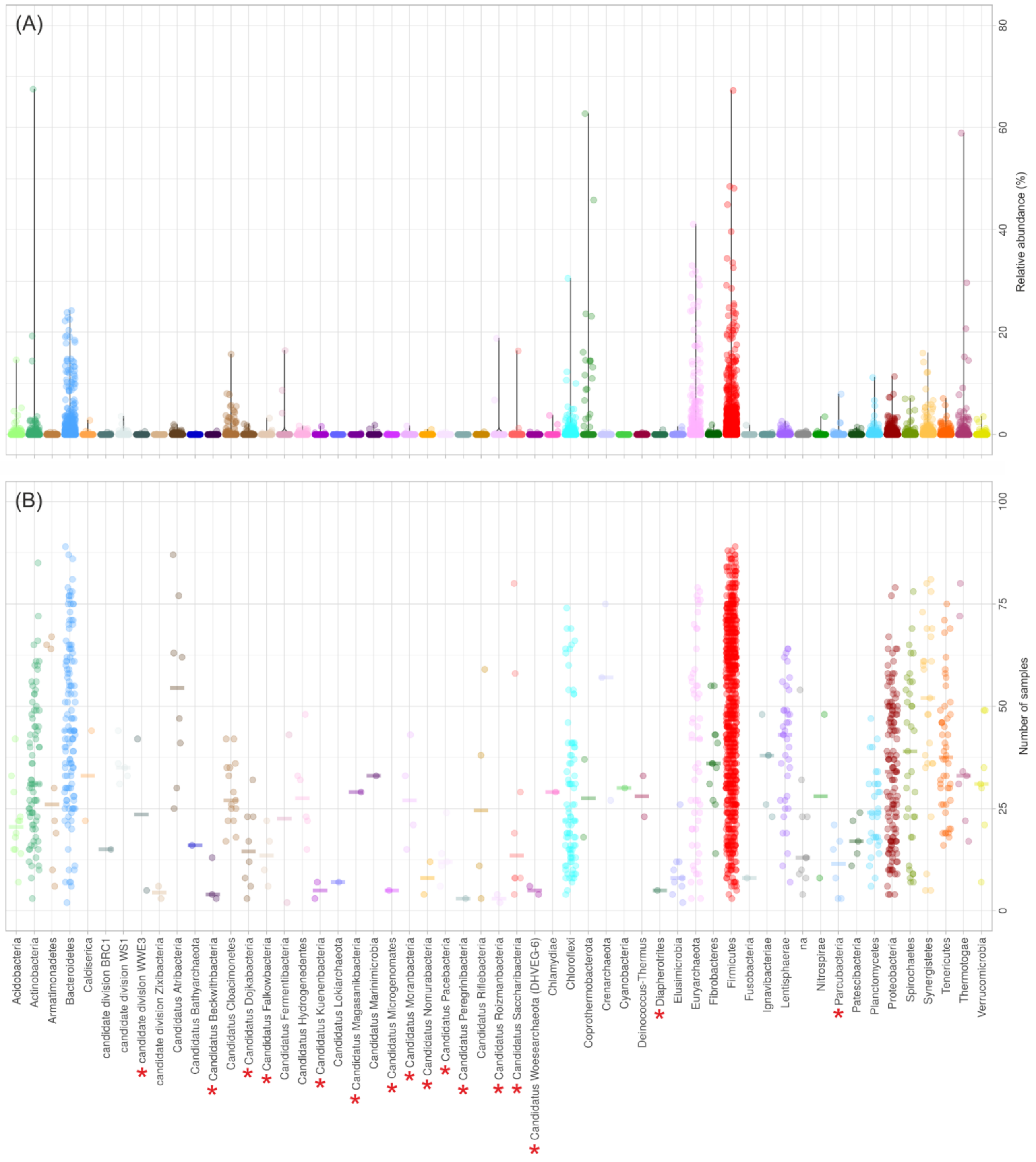


**Additional Fig. S4.** (A) Dots represent MAGs relative abundance in the samples examined. Dots are colored according to the taxonomic assignment of the MAGs at phylum level. (B) Number of samples where each MAG was identified at relative abundance higher than 0.001%. Each dot is representative of a single MAG and the number of samples where it was identified is reported in y axes (CPR are marked with asterisks).
